# Outer Membrane Vesicles Mediated Horizontal Transfer of an Aerobic Denitrification Gene between *Escherichia coli*

**DOI:** 10.1101/835694

**Authors:** Yang Luo, Jiahui Miao, Weichuan Qiao

## Abstract

Bacterial genetic material can be horizontally transferred between microorganisms via outer membrane vesicles (OMVs) released by bacteria. Up to now, the application of vesicle-mediated horizontal transfer of “degrading genes” in environmental remediation has not been reported. In this study, the *nirS* gene from an aerobic denitrification bacterium, *Pseudomonas stutzeri*, was enclosed in a pET28a plasmid, transformed into *Escherichia coli* (*E. coli*) DH5α and expressed in *E. coli* BL21. The *E. coli* DH5α released OMVs containing the recombination plasmid pET28a–*nirS*. Moreover, the amount of released OMVs-protein and DNA in OMVs increase as heavy metal concentrations and temperature increased. When compared with the free pET28a–*nirS* plasmid’s inability to transform, *nirS* in OMVs could be transferred into *E. coli* BL21 with the transformation frequency of 2.76×10^6^ CFU/g when the dosage of OMVs was 200 µg under natural conditions, and *nirS* could express successfully in recipient bacteria. Furthermore, the recipient bacteria that received OMVs could produce 18.16 U ml^-1^ activity of nitrite reductase. Vesicle-mediated HGT of aerobic denitrification genes provides a novel bioaugmentation technology of nitrogen removal.

**Importance:** Previous studies have reported that bacterial genetic material can be horizontally transferred between microorganisms via outer membrane vesicles(OMVs) released by bacteria. However, the application of vesicle-mediated horizontal transfer of “degrading genes” in environmental remediation has not been reported. In this study, we found that OMVs could mediate horizontal transfer of pET28a–*nirS* plasmid between E. coli under natural condition. The transformation frequency reached to 2.76×10^6^, which was higher than that of the free plasmid. Vesicle-mediated HGT of aerobic denitrification genes provides a novel bioaugmentation technology of nitrogen removal.

## Introduction

High nitrogen concentrations in water result in water eutrophication and pollution. Traditional bio-treatment processes for nitrogen removal involve autotrophic and heterotrophic denitrification under aerobic and anoxic conditions, respectively (1). Because of their different oxygen requirements, these two steps are separated spatially and temporally. Recently, more researchers have focused on nitrogen removal using aerobic denitrification bacteria. Unlike traditional anaerobic denitrification mechanisms, aerobic denitrification occurs via co-respiration or co-metabolism of O_2_ and NO_3_^-^. Additionally, nitrification and denitrification can occur in the same aerobic denitrification system (2, 3). Nitrate reductase (*napA*), nitrite reductase (*nirS*), nitric oxide reductase (*norB*) and nitrous oxide reductase (*nosZ*) are four key enzymes in the aerobic denitrification process of aerobic denitrifying bacteria. The aerobic denitrification bacteria or microbial consortium can be added into wastewater for promoting the removal efficiency of nitrogen (4). In addition, gene bioaugmentation is considered to be a promising biological enhancement process, which also promotes pollutant removal by horizontal gene transfer (HGT) (5).

HGT is common and important in microorganisms because it enables the bacteria to accept exogenous functional gene fragments to obtain new metabolic functions and then present a new ecological phenotype (6, 7). Artificially constructed plasmids with degradation genes to accelerate the horizontal transfer of these genes in contaminated areas have become a research hotspot (8, 9). HGT provides a new idea for bioaugmentation technology in the field of environmental remediation; indigenous microorganisms in a contaminated system obtain degradation genes from self-apoptosis of recombinantly engineered bacteria or the natural release of bacteria to improve the pollutants’ degradation efficiency. However, free-DNA in the environment is easily degraded by DNase I enzyme, and high temperatures cause the DNA to break down naturally.

In the past 30 years, studies have found that a wide variety of Gram-negative bacteria (i.e., *Escherichia coli* (10), *Bacteroides thetaiotaomicron* (11), *Pseudomonas aeruginosa* (12), *Piscirickettsia salmonis* (13), *Acetobacter pasteurianus* (14), *Acinetobacter baumannii* (15) and *Helicobacter pylori* (16) can release outer membrane vesicles (OMVs), a kind of spherical nanometer-sized proteolipids. The analysis of vesicle components revealed vesicles contained outer membrane proteins (i.e., OM-anchored lipoproteins, OM phospholipids and LPS), inner membrane proteins, periplasmic components (i.e., periplasmic protein and hydrolase), signaling molecules, virulence factors, DNAs and RNAs (17–22). Most research indicates that vesicles play an important role in bacterial life activities. Vesicles can be used as carriers for transporting signaling molecules (20, 23, 24), virulence factors(25, 26) and resistance genes (11, 27) to other bacteria. Reports suggest that genes can be transferred via OMVs. S. Fulsundar et al. (28) found that exposure to OMVs isolated from plasmid-containing donor cells results in HGT to *A. baylyi* and *E. coli* at transfer frequencies ranging from 10^-6^ to 10^-8^, with transfer efficiencies of approximately 10^3^ and 10^2^ per μg of vesicular DNA, respectively. C. Rumbo et al. (29) provided evidence that carbapenem resistance genes OXA-24 were delivered to surrounding *A. baumannii* bacterial via vesicles. In addition, chromosomal DNA presented in OMVs was found to transfer between *P. gingivalis* at a frequency of 1.9×10^-7^ (30). S. Yaron et al. (25) showed vesicle-mediated transfer of plasmid and phage DNA from *E. coli* O157:H7 to *E. coli* JM109 and phage DNA to *S. enterica* serovar Enteritidis ATCC 13076. H. X. Chiura et al. (31) revealed that the highest gene transfer frequencies were up to 1.04×10^2^ via vesicles harvested from seawater.

However, research on horizontal transfer of genes via vesicles focuses mainly on the horizontal transfer of antibiotic genes via vesicles on global bacterial resistance (11, 28, 32) or vesicle-mediated virulence genes in bacterial pathogenesis (25). There are few studies on the horizontal transfer of “degradation genes” via vesicles in repairing environmental pollution. With the rapid development of chemical, petroleum, synthetic ammonia and cooking industries, a slew of nitrogen-containing substances is discharged into bodies of water, resulting in eutrophication.

In this study, the nirS gene cloned from an aerobic denitrification bacteria, *Pseudomonas stutzeri* was inserted into a pET28a plasmid, and thereby transformed to *E. coli* DH5α. We evaluated the characteristic of OMVs released from the recombinant strain and studied the vesicle-mediated HGT of the *nirS* gene to enhance the denitrification efficiency of microorganisms in nitrogen-containing wastewater. We hope to explore a novel bioaugmentation technology using HGT for biological denitrification technology.

## Results

### Cloning and analysis of the EGFP and full-length nirS

EGFP, the green fluorescent protein gene, was cloned from a laboratory preservation solution, and the sequencing results were consistent with the NCBI database (GenBank Accession: AFA52650.1). The gene *nirS* is one of the four aerobic denitrification genes, which can convert NO_2_^-^ to NO by a reduction reaction. The full-length DNA (1713 bp) of *nirS* was amplified from *P. stutzeri* by PCR and deposited into GenBank under accession number MN199166. The open reading frame (ORF) encodes a protein of 570 amino acids. The molecular weight of *nirS* was approximately 63 kDa and was consistent with the estimated molecular weight of the enzyme. The nucleotide and amino acid sequences of the full-length DNA are shown in Fig. S1. The sequence similarity of the nirS gene from *P. stutzeri* BNCC 139708 with the homologous strains *P. stutzeri* CCUG 29243, *P. stutzeri* CGMCC 1.1803, *P. stutzeri* R2A2 and *P. stutzeri* 19SMN4 was 99.47%, 98.07%, 95.80% and 94.83%, respectively. The similarity of their amino acid sequences was 100%, 99.79%, 95.80% and 94.92%, respectively. The secondary structures of their proteins encoding *nirS* were similar, and there was no difference except for the N-terminus of *P. stutzeri* 19 SMN 4 (Fig. S2). The proportions of alpha-helix, extended strand, beta turn and random coil in the five *P. stutzeri* were approximately similar. These results indicated that *P. stutzeri* BNCC 139708 was highly homologous with other four species of *P. stutzeri*, and the *nirS* gene was conservative in the evolution process.

The analysis of signal peptide sequence in the *nirS* showed that the C and Y value were the highest and the S value was steep at the 24th amino acid where was the predicted signal peptide cutting site (Fig. S3). The 1-23 amino acid was the position of the nitrite reductase signal peptide. The number of transmembrane helix amino acid residues in the *nirS* was 2.83622, and the amount of transmembrane helix amino acid in the first 60 amino acids was 2.83343. Fig. S4 showed that there was a transmembrane helix signal in the first 100 amino acids which was the signal peptide sequence in the *nirS*. So the protein encoding the *nirS* was not a transmembrane protein. The *nirS* gene could be constructed into a vector for prokaryotic expression.

### Construction and expression of pET28a-nirS-EGFP and pET28a-nirS plasmids

In this study, the plasimids harboring *napA*, *norB*, *nirS* and *nosZ* gene were constructed, respectively. However, only the *nirS* gene could be expressed in *E. coli* BL21; therefore, we selected the plasmid harboring *nirS* as the object gene of HGT. The construction of pET28a-*nirS*-EGFP and pET28a-*nirS* plasmids is shown in Fig. 1a and Fig. S6. The two plasmids were transformed and expressed in *E. coli* BL21(DE3). As shown in Fig. 1b, *E. coli* BL21(DE3), containing the recombinant plasmid pET28a-*nirS*, was induced by 0.2 mM IPTG to express soluble 63 kDa. The optimal induced expression condition of pET28a-*nirS* was 16 °C at 110 rpm for 16 h. For *E. coli* BL21(DE3) containing the recombinant plasmid pET28a-*nirS*-EGFP, the SDS-PAGE after IPTG induction showed a band at approximately 89 kDa, corresponding to the expected size of the fusion protein (Fig. 1c, lane 2). For BL21(DE3) cells transformed with empty pET28a, SDS-PAGE after IPTG induction showed no band (Fig. 1c, lane 1). Furthermore, western blotting results showed there were obvious bands at the corresponding positions, indicating that *E. coli* BL21(DE3) successfully expressed the fusion protein (Fig. 1d).

**Figure 1.**
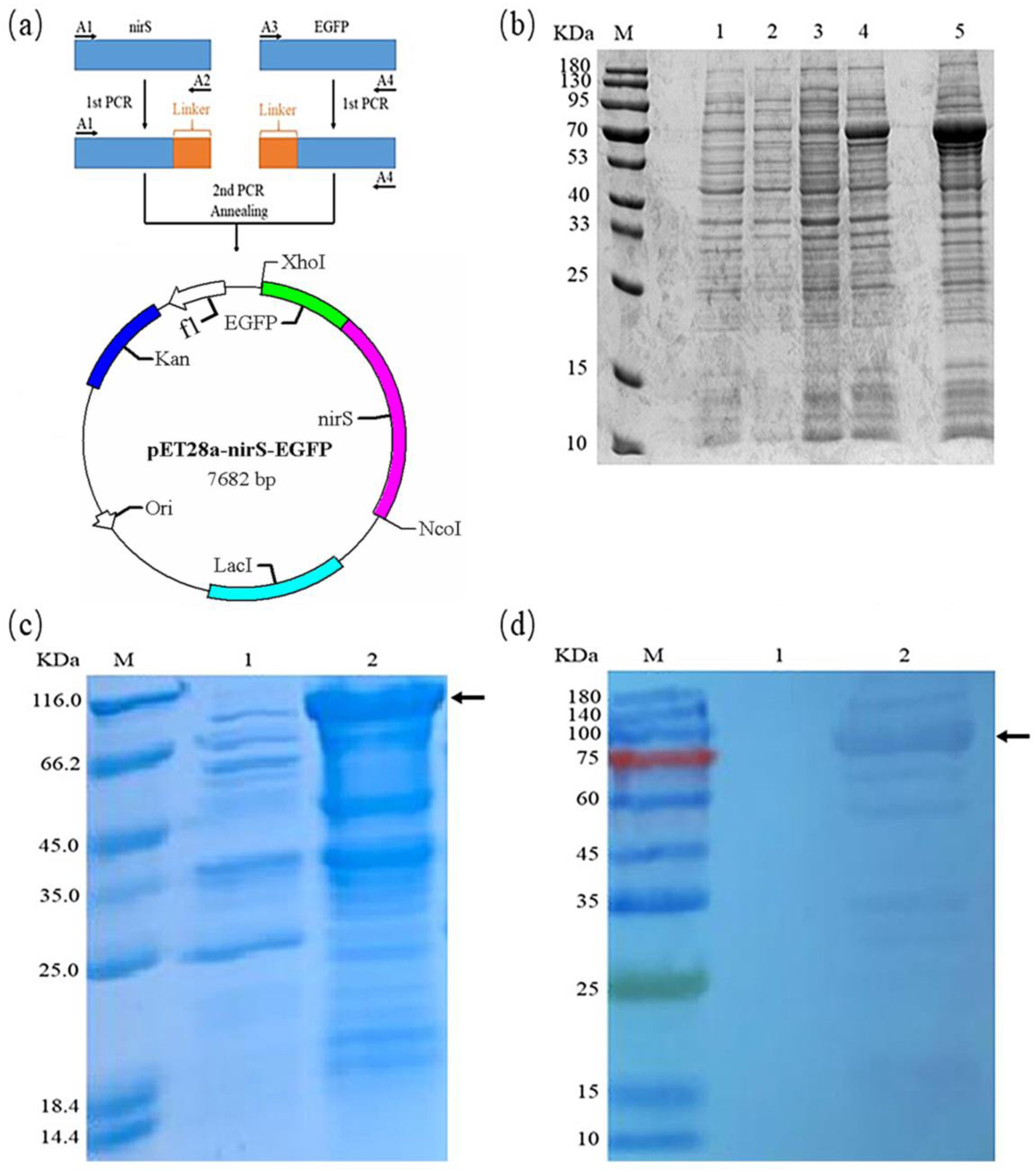
(a) Schematic representation of the construction of the recombinant vector, pET28a-*nirS*-EGFP. Arrows indicate primers used. (b) SDS-PAGE analysis of induced *E. coli* BL21(DE3) containing recombinant plasmid pET28a-*nirS* under different conditions. M: protein marker; Lane 1: induced expression of empty vector pET28a; Lane 2: 37 °C, 150 rpm; Lane 3: 28 °C, 130 rpm; Lane 4: 22 °C, 120 rpm; Lane 5: 16 °C, 110 rpm. (c) SDS-PAGE analysis of *nirS*-EGFP fusion protein. M: protein marker; Lane 1: lysates of bacteria transformed with empty pET28a under IPTG induction. Lane 2, *nirS*-EGFP fusion protein purified by immobilized metal affinity chromatography. Arrows indicate the target protein. (d) Western blotting of *nirS*-EGFP fusion protein using an anti-His6 tag. M: protein marker; Lane 1: lysates of bacteria transformed with empty pET28a. Lane 2, induced expression of *nirS*-EGFP fusion protein. Arrows indicate the target protein.

### OMVs released from E. coli harboring the plasmid pET28a-nirS

In this study, we transformed the plasmid pET28a-*nirS* into *E. coli* DH5α and observed if the transformed *E. coli* could release OMVs. As shown in Fig. 2a and b, *E. coli* DH5α harboring the plasmid could release OMVs. OMVs production by *E. coli* strain was confirmed by transmission electron microscopy, indicating that *E. coli* actively released OMVs without any external stimulation during its growth. OMVs were isolated from bacteria in late stationary phase by an ultra-high-speed freezing centrifuge. Fig.2a indicates the process of OMVs released from the outer membrane of *E. coli*. OMVs have a spherical structure with a bilayer membrane, with an average diameter of 25 nm. The solution containing OMVs were observed with a microscope and coated on a plate with Luria-Bertani agar (LA) medium. There were no bacteria found under the microscope and no bacterial growth on the plate overnight, indicating that OMVs extracted in this study had high purity.

**Figure 2.**
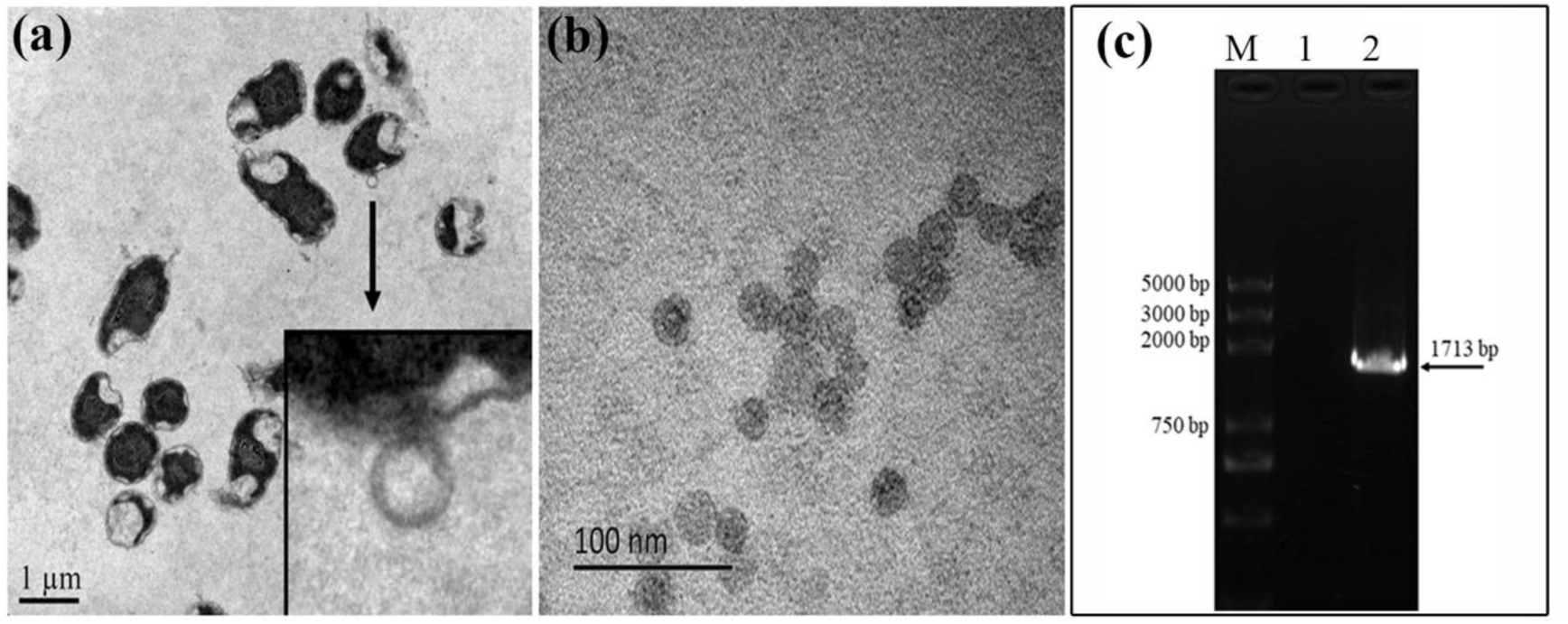
TEM of OMVs released from *E. coli* DH5α (a, b) and PCR amplification products of aerobic nitrification genes in the OMVs (c). Lane 1, Gel electrophoresis band of supernatants removed any *E. coli* DH5α and OMVs; lane 2, Gel electrophoresis band of *nirS* isolated from OMVs from *E. coli* DH5α.

DNA in the OMVs from *E. coli*-*nirS* was amplified by PCR using specific primers for the *nirS* gene. As shown in Fig. 2c, there was a band of *nirS* genes, showing that OMVs carried the pET28a-*nirS* plasmid, but the typical *nirS* band was not detected in the supernatant removed from *E. coli*-*nirS* and OMVs. SDS-PAGE analysis of protein concentration of OMVs from *E. coli* DH5α under different stress conditions is shown in Fig. S5.

### Effect of environmental stresses on vesiculation and inclusion content in OMVs

To evaluate the effect of environmental stresses on OMVs released from bacteria, heavy metals (Cu^2+^ 25 mg l^-1^, 50 mg l^-1^; Cd^2+^ 5 mg l^-1^) were added to the culture medium to stimulate the release of OMVs from *E.coli*. As shown in Figs. 3 and 4, the concentration of inclusions (protein and DNA) in OMVs and diameter of OMVs increased with increasing concentrations of Cu^2+^. In addition, OMVs’ diameter increased when the concentration of Cd^2+^ increased to 5 mg/L but decreased when Cd^2+^ increased to 25 mg l^-1^. The results showed that different metals had different effects on the release of OMVs. In contrast, temperature increases from 25 °C to 37 °C caused a decrease in OMVs’ diameter while increasing the concentrations of protein and DNA in OMVs from *E.coli*.

**Figure 3.**
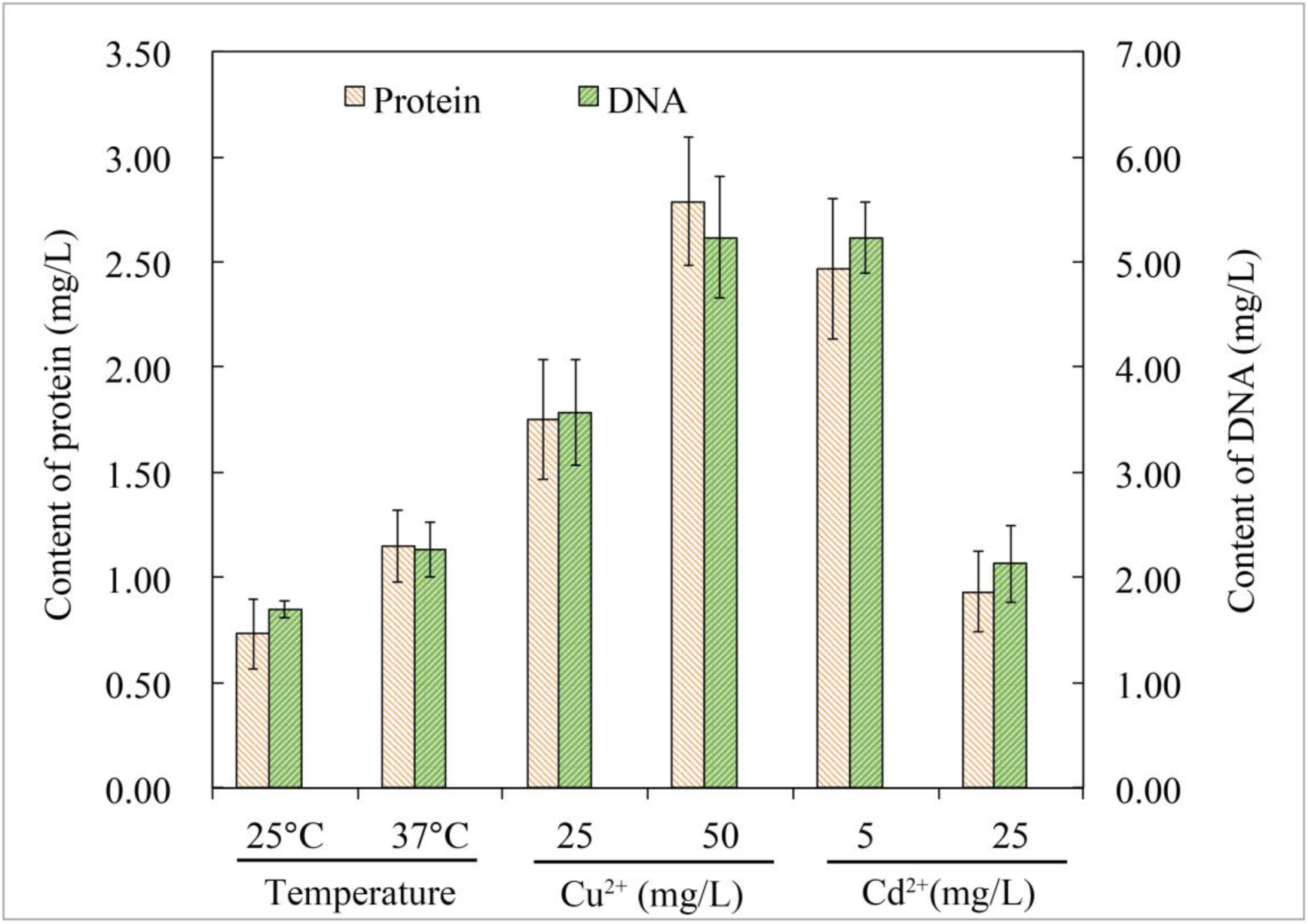
Protein and vesicular DNA concentrations from OMVs isolated from *E. coli* under different stress conditions.

**Figure 4.**
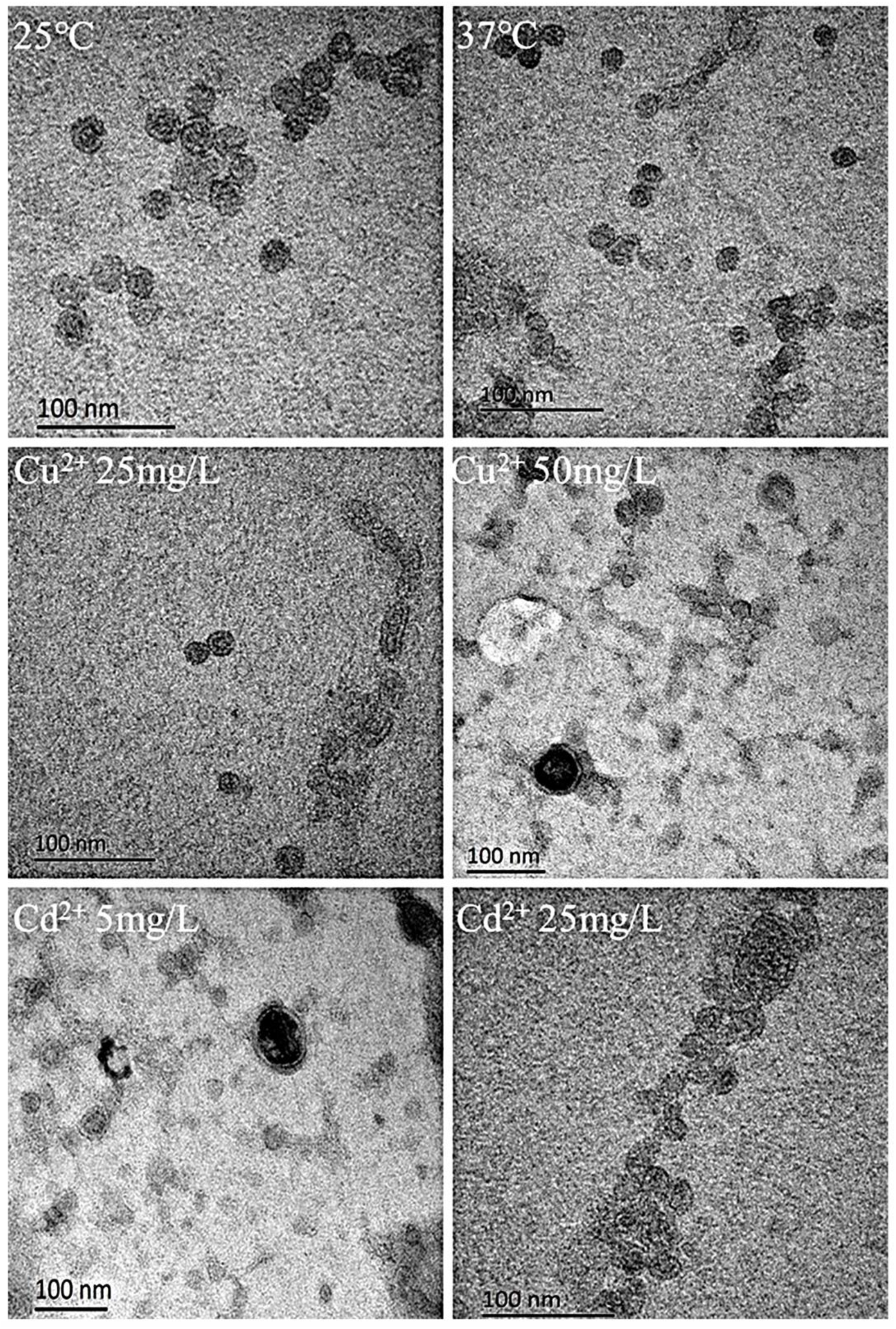
OMVs released by *E. coli* under different stresses.

### Horizontal transfer of the aerobic denitrification gene (nirS)

In this study, the membrane of OMVs released from *E. coli*-*nirS*/EGFP was fluorescently stained with CM-DiI dye (orange color) (Fig. 5, upper panel), and *E. coli* BL21 was fluorescently stained with DiO dye (green color). The stained OMVs and *E. coli* BL21 were incubated together to observe vesicle-mediated horizontal transfer of *nirS* from *E. coli* DH5α to *E. coli* BL21. After 1 h incubation, the receptor *E. coli* BL21 simultaneously emitted green and orange fluorescence under a fluorescence microscope (Fig. 5, lower panels). OMVs and *E. coli* BL21 were observed as light green under confocal laser microscopy (LCSM), indicating that free OMVs could directly contact and adsorb on the cell membrane of the receptor *E. coli* BL21 cell. After culturing OMVs released from *E. coli*-*nirS*/EGFP added to a culture medium containing *E. coli* BL21, all the bacterial liquid was uniformly coated on an LA plate with kanamycin resistance. Individual clones were selected for culture and induced with IPTG after which we observed green fluorescent protein expression in the *E. coli* BL21 strain using LCSM (Fig. 6). The results showed that *E. coli* BL21 could produce green fluorescence, suggesting that the aerobic denitrification gene (*nirS*) had been horizontally transferred to *E. coli* BL21 via OMVs and successfully expressed in *E. coli* BL21. The SDS-PAGE analysis of protein product from the receptor *E. coli* BL21 showed significant expression of the *nirS* gene. The expressed nitrite reductase was detected under a different induction temperature; moreover, the temperature greatly affected nitrite reductase activity. In addition, reductase activity reached to 18.16 U ml^-1^ under an optimal induced temperature of 22 °C as shown in Fig. 7.

**Figure 5.**
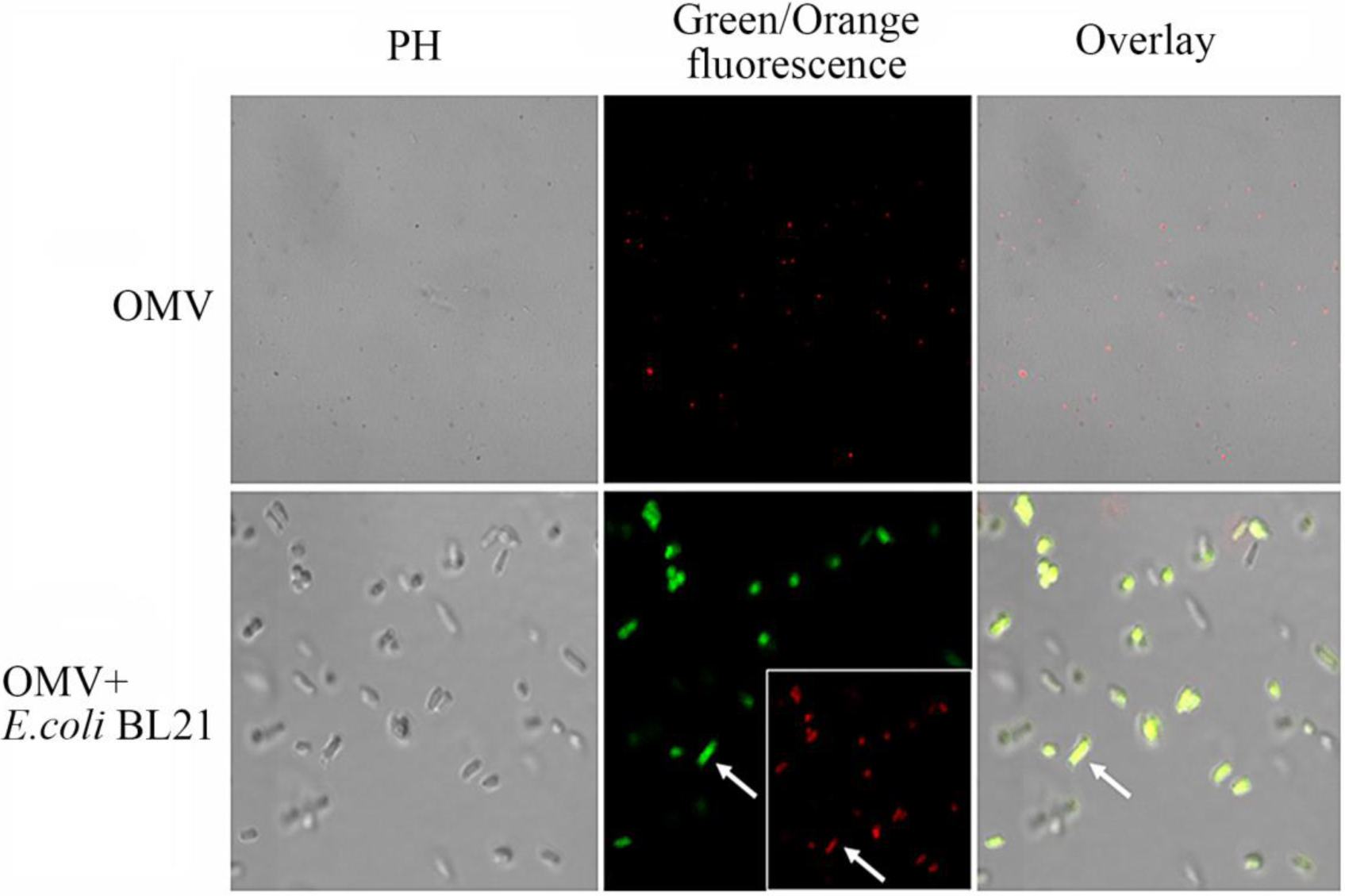
Recombinant plasmid *nirS*-EGFP was delivered via OMVs to non-producing cells. (PH, phase contrast images; green/orange fluorescence, green/orange fluorescence images.)

**Figure 6.**
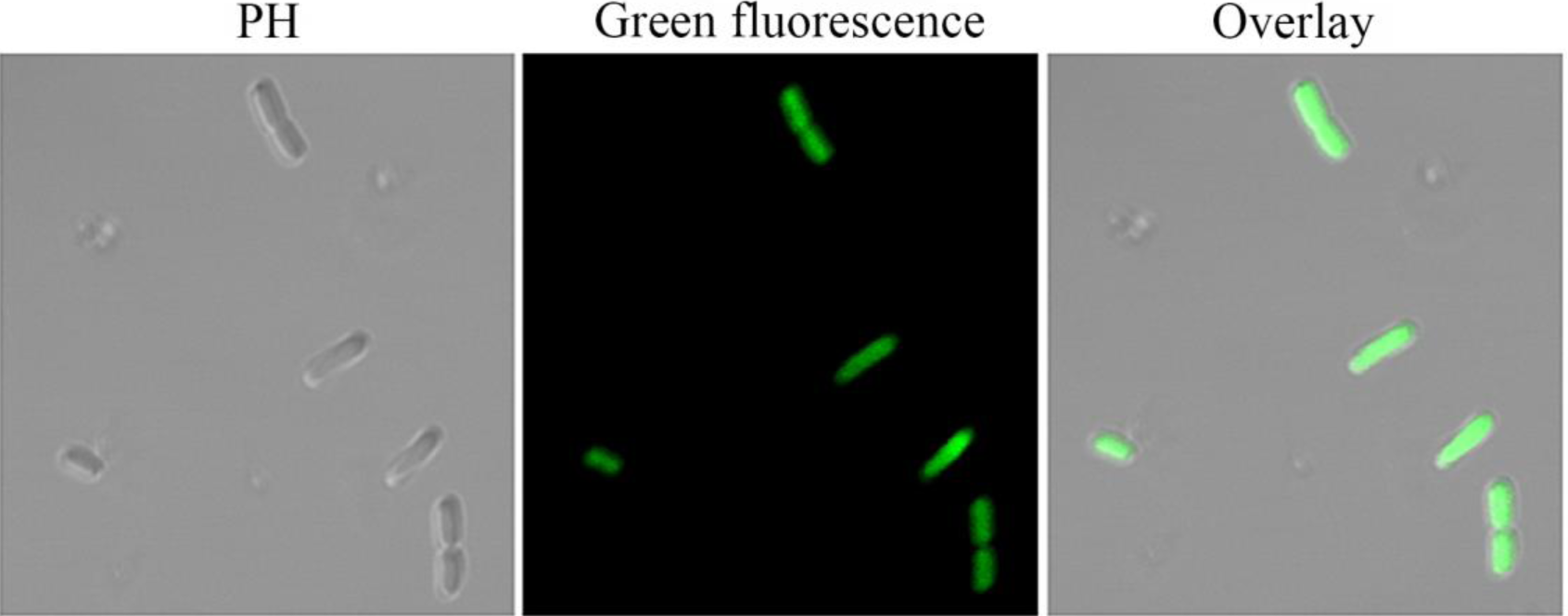
The LCSM images of *E. coli* BL21 induced by IPTG. PH: phase contrast image; Green fluorescence: green fluorescence image; Overlay: image of phase contrast image and green fluorescence image superimposed.

**Figure 7.**
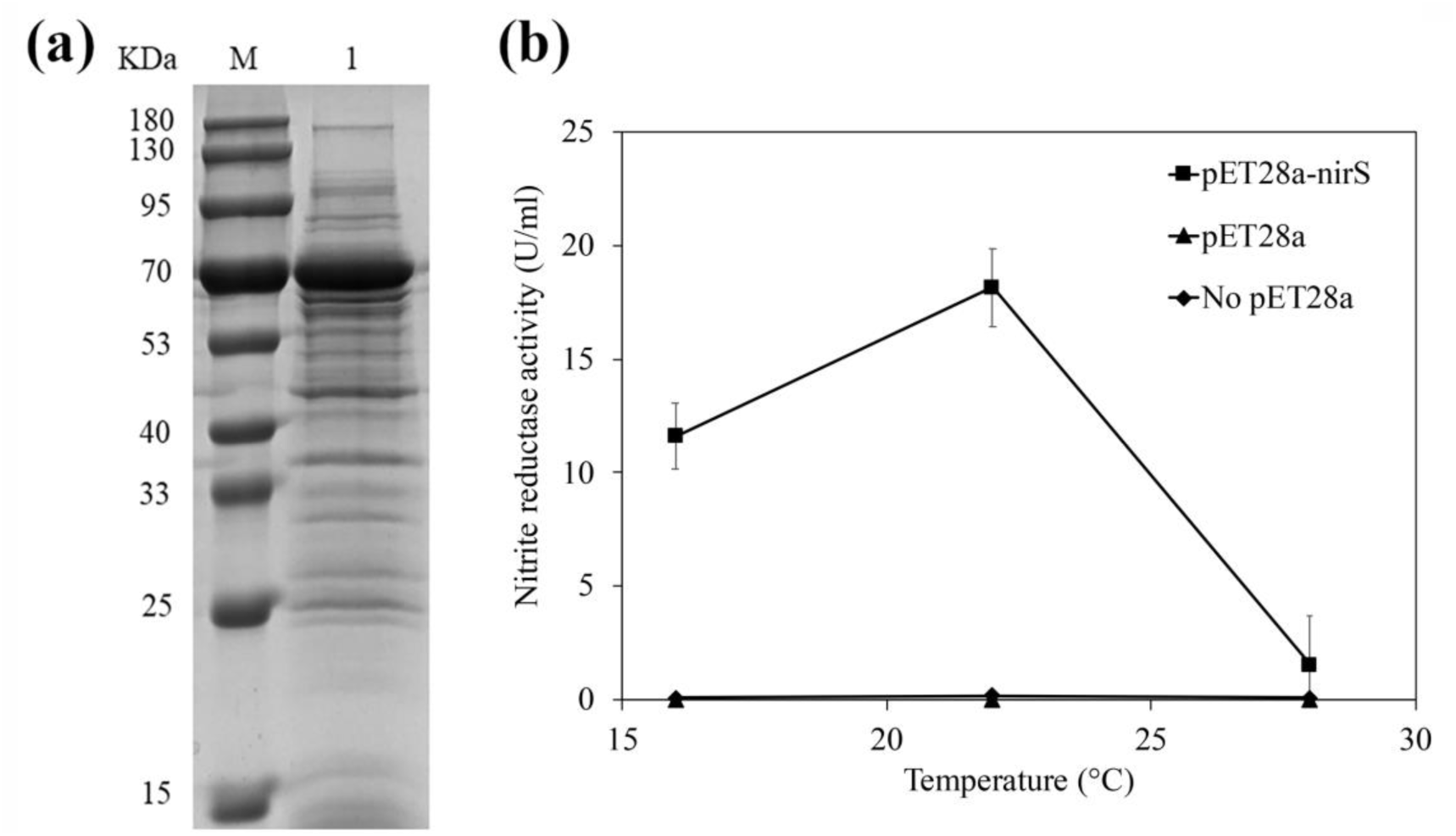
SDS-PAGE analysis and activity of nitrite reductase expressed by *E. coli* BL21. (a) M: protein marker; Lane 1: protein product. (b) Nitrite reductase activity of *E. coli* BL21 transformed via OMVs under different temperatures.

### Transformation frequency of OMVs

Transformation experiments were successful in *E. coli* BL21 via OMVs isolated from *E. coli*-*nirS* with different amounts (50, 100 and 200 µg of protein). As shown in Fig.8, when the dosage of OMVs was 200 µg, the transformation frequency of *nirS* reached to 2.76×10^6^ CFU/g; moreover, it increased with the dosage of OMVs. No transformants were obtained when free pET28a-*nirS* plasmid (20 ng) was incubated with *E. coli* BL21, indicating that free plasmid DNA could not be transformed into recipient cells without OMVs as a “protective shell” or with any treatment of the recipient bacteria (preparation of competent cells through heat-shock during transformation). The pET28a-*nirS* plasmid protected by OMVs successfully transferred to recipient strains in a dose-dependent manner.

**Figure 8.**
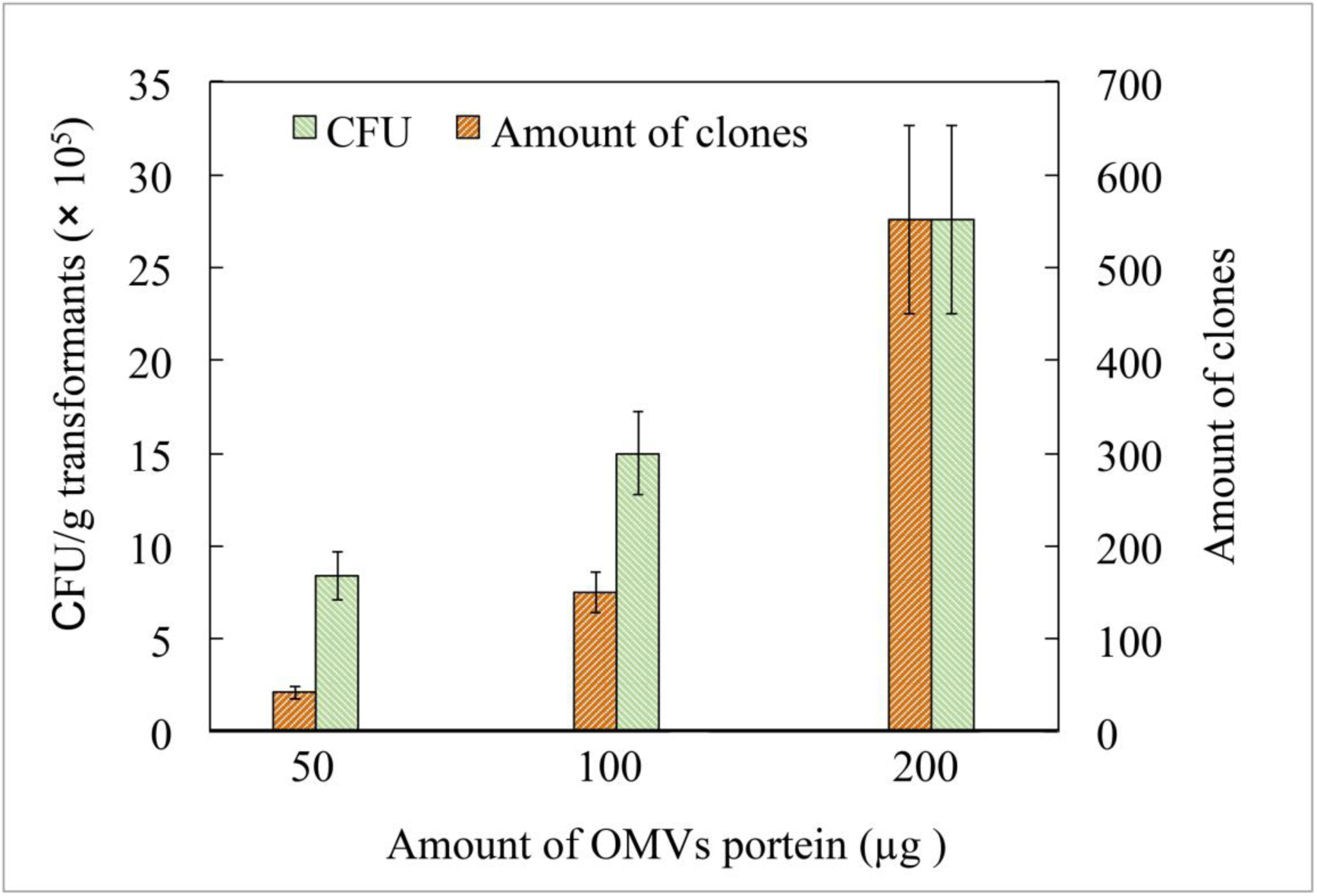
Transformation frequency of the *nirS* gene by OMVs.

## Discussion

Previous studies have reported that vesicles released from the bacterial outer membrane are loaded with many bacterial contents. Different types of nucleic acids (i.e., chromosomes, plasmids, DNA and RNA) were found in OMVs released from Gram-negative bacteria, as shown in Table 1. In this study, the gene *nirS* was also detected in vesicles released by recombinant *E. coli*-*nirS*. Moreover, the amount of released vesicles containing DNA was affected by temperature and heavy metals.

**Table 1.** Types of genetic material in vesicles released by different bacteria

| Gene | Bacterial species | Reference |
| --- | --- | --- |
| DNA |  |  |
| Chromosomal DNA | *E. coli*, *Shewanella vesiculosa*, *Thermococcales*, *P. aeruginosa* | (48-50) |
| Plasmid DNA | *E. coli*, *B. burgdorferi*, *A. baylyi*, *P. aeruginosa* | (22, 28, 47, 51) |
| RNA |  |  |
| mRNA | *E. coli*, *P. aeruginosa*, | (21, 52) |
| tRNA | *E. coli*, *P. aeruginosa*, *B. burgdorferi* | (21, 22, 52, 53) |
| rRNA | *E. coli* | (22, 52, 53) |
| miRNA | *P. gingivalis*, *Treponema denticola* | (54) |
| sRNA | *P. aeruginosa*, *B. burgdorferi* | (21, 22, 55) |

Reports indicate that different stress conditions, including antibiotics (28), temperature (10), ultraviolet irradiation (28, 33), virulence factors (34), SOS response (35), oxygen stress (36), nutritional deprivation (28) and physical pressure (34), can affect the number of vesicles released from bacteria. In this study, the release of OMVs was stimulated by adding heavy metals to the environment when the bacteria growth rate was unrestricted. The results showed that the release of OMVs increased with the increase of heavy metal toxicity. The Tol–Pal, an inner membrane protein systems of gram-negative bacteria, interacts with peptidoglycan (PG) to maintain cell membrane stability. Studies have shown that disruption of the Tol-Pal system or lack of interaction in the outer membrane protein-peptidoglycan and the inner membrane protein-peptidoglycan induces membrane instability, causing cells to increase vesicle biogenesis to maintain membrane stability (37, 38). Heavy metal toxicity may cause damage to the cell membrane, causing local deletion of the Tol-Pal system or a decrease in the amount of membrane protein and resulting in the release of a large amount of OMVs by *E. coli*. However, when the concentration of Cd^2+^ increased to 25 mg l^-1^, the amount of OMVs released by bacteria decreased. This may be due to bacterial growth rate inhibition under the stimulation of high heavy metal concentrations, reducing the amount of OMVs released by cells in the early stage of bacterial growth. Recent studies have also shown that high growth temperatures can increase the number of vesicles produced. In this study, when the culture temperature of the *E. coli* increased from 25 °C to 37 °C, the number of produced OMVs increased. The temperature stress increases the accumulation of misfolded proteins, which will be transported out from bacteria by promoting the formation of OMVs to maintain normal cell physiological activity (10).

In this study, when compared with the control, DNA content increased with the increase of heavy metal toxicity and temperature without affecting the growth rate of bacteria. In addition, the diameter of OMVs released by *E. coli* increased under temperature and heavy metal stress, possibly due to serious distortions in cell membranes. Moreover, it was easier for larger OMVs to load intracellular substances, resulting in increased DNA content in OMVs.

Horizontal transfer of specific degradation genes not only improves the survival rate of microorganisms in the system but also enhances the biodegradation efficiency of pollutants (39–41). Many studies have shown that vesicles can accelerate the flow of genes between microorganisms. It is valuable for ecological environment restoration efforts to study the horizontal transfer efficiency of pollutant degradation genes between microorganisms under vesicle protection. In this study, it was not only confirmed that OMVs from *E. coli* were loaded with aerobic denitrification genes but also found that the aerobic denitrification gene *nirS* could be successfully transferred to *E. coli* BL21 via OMVs and expressed in *E. coli* BL21. This indicates that OMVs-mediated horizontal transfer of “degradation genes” promotes the degrading ability of microorganisms and plays an active role in pollution remediation.

Vesicles in the environment transfer DNA into the recipient cell by fusing with the outer membrane (23), or entire vesicles are absorbed into receptor cells by dynamin-dependent endocytosis (26, 42). These pathways are equally applicable to the transport of virulence factors between bacteria and vesicles. In this study, we labeled OMVs and *E. coli*-*nirS* with fluorescence and found that free OMVs could be directly adsorbed on the cell membrane of *E. coli* BL21. Previously, it was reported that human intestinal epithelial cells (Caco-2), brain microvascular endothelial cells (HBMEC) and glomerular endothelial cells (HRGEC) could ingest vesicles labeled with rhodamine isothiocyanate B-R18 by endocytosis of kinase protein (26). In the present study, it was also observed that OMVs could be internalized into cells with no functional hindrance; OMVs fused with the receptor after contacting receptor cells and released substances into the recipient cytoplasm. The discovery of this process provides a practical basis for enhancing genetic material exchange mediated by OMVs in bacterial communities.

Studies of these bacterial OMVs support a common role of vesicles on easy exchange of genetic material between bacteria (29, 43, 44), indicating that vesicle-mediated HGT could enhance gene fluidity among species and improve the survival rate of sensitive bacteria in the environment. The mechanism can be applied to the field of environmental remediation. Bacteria in a polluted environment can obtain degradation genes by vesicles to improve the efficiency of pollutant removal. In this study, after adding OMVs loaded with pET28a-*nirS*-EGFP recombinant plasmid into liquid medium containing *E. coli* BL21, we discovered that *E. coli* BL21 could express nitrite reductase and green fluorescent protein, showing that bacteria obtained the ability to reduce nitrite. Moreover, the process of DNA transformation mediated by OMVs was dose-dependent because the frequency of HGT increased with the increase of OMVs.

## Materials and methods

### Bacteria and growth conditions

*Pseudomonas stutzeri* (BNCC 139708) was purchased from BeNA Culture Collection, China. *P. stutzeri* grew in beef-extract peptone medium (0.3% beef cream, 1% Peptone, 0.5% NaCl) at 30°C on a shaker at 150 rpm. *E. coli* DH5α and *E. coli* BL21 (DE3) were purchased from Sagon Biotech, Shanghai, China, which grew in Luria-Bertani broth (LB) medium (1% tryptone, 0.5% yeast extract, 1% NaCl) at 35°C on a shaker at 150 rpm or LA medium (1% tryptone, 0.5% yeast extract, 1% NaCl and 2% agar in an incubator) at 35 °C with Kanamycin (50mg l^-1^) for selecting positive recombinants.

### Molecular cloning of full-length nirS and EGFP

The DNAMAN software was used to compare multiple *nirS* genes of *P. stutzeri* on the NCBI website to design homologous primers. The full-length *nirS* was generated from *P. stutzeri* by PCR using homologous primers N1 and N2 (Table S2). The procedures were as follows: 94 °C for 4 min followed by 35 cycles (94 °C for 60 s, 62 °C for 40 s and 72 °C for 10 min). The full-length EGFP was generated by PCR with primers EGFP-P1 and EGFP-P2 (Table S2). The procedures were as follows: PCR was performed for 30 cycles at 94 °C for 30 s, 58 °C for 30 s and 72 °C for 60 s. TA cloning was carried out after purifying the PCR products by cutting gel recovery, and five recombinant plasmids were screened and sequenced by Shanghai Biotechnology Service Company. The sequence of the *nirS* was submitted on NCBI website (http://www.ncbi.nlm.nih.gov) which can be used for homology analysis. The secondary protein structure of the *nirS* was analyzed on SOPMA website (https://npsa-prabi.ibcp.fr). The signal peptide sequence of the gene was analyzed by Detaibio bioinformatics tools (http://www.detaibio.com/tools/). The presence and Location of transmembrane region in the *nirS* sequence was predicted by TMHMM2.0 (http://www.cbs.dtu.dk).

### Construction, expression and purification of nirS-EGFP and nirS

We used overlap PCR to generate the *nirS*-EGFP fusion gene. The following four primers, designed based on the sequences of EGFP and *nirS*, were used (Table S2). The *nirS* gene was cloned by PCR using primers A1 and A2 with an extra NcoI recognition site at its 5′ end. The procedures were as follows: 95 °C for 3 min followed by 35 cycles (95 °C for 22 s, 60 °C for 20 s and 72 °C for 50 s). The primers A3 and A4 were used to amplify the DNA sequence of EGFP, which contains an XhoI recognition site at its 3′ end. PCR was performed for 30 cycles of 95 °C for 22 s, 60 °C for 20 s and 72 °C for 50 s. Products of the first PCR round were gel purified and subjected to a second round of PCR using A1 and A4 primers to generate a product encoding *nirS*-EGFP, containing a linker-encoding sequence between the two domains. Amplification was performed for 30 cycles of denaturation for 22 s at 95 °C, annealing for 20 s at 60 °C and extension for 75 s at 72°C. The *nirS* and *nirS*-EGFP PCR products were digested with BamHI and Hind III or NcoI and XhoI, respectively, and ligated into the pET28a plasmid at the corresponding restriction sites. The construction of plasmids pET28a-*nirS* and pET28a-*nirS*-EGFP is shown in Figs. S3 and S4. Recombinant plasmid pET28a-*nirS* and pET28a-*nirS*-EGFP were transformed into *E. coli* DH5α competent cells for propagation of recombinant plasmids, which was named *E. coli*-*nirS* or *E. coli*-*nirS*/EGFP.

The recombinant plasmids pET28a-*nirS* and pET28a-*nirS*-EGFP were transformed into competent *E. coli* BL21 (DE3) cells to express the fusion protein. LB medium of 50 ml (50 μg ml^-1^ kanamycin) was inoculated (1:50) with *E. coli* BL21 with recombinant plasmid pET28a-*nirS* suspension and grown at 37 °C at 220 rpm until reaching an optical density at 600 nm (OD600) of 0.4–0.6. Then, expression was induced by adding 0.5 mM isopropyl-D-1-thiogalactopyranoside (IPTG), and the cultures were incubated for 16 h at 16 °C at 150 rpm. Bacteria were harvested by centrifugation at 8000×g for 25 min at 4 °C. Pellets were treated with sonication on ice for a 4 s pulse with an intervening 4 s pause until cells were completely lysed. Lysates were centrifuged at 15,000×g for 20 min at 4 °C. The supernatants were collected to analyze them on polyacrylamide gel electrophoresis (PAGE).

To produce soluble pET28a-*nirS*-EGFP proteins, the following induction scheme was established: final IPTG concentration, 0.2 mM; induction temperature, 16 °C; total induction duration, 16 h; shaking speed, 110 rpm. After induction, bacteria were harvested by centrifugation at 8000×g for 25 min at 4 °C. Pellets were treated with sonication on ice for a 4 s pulse with an intervening 4 s pause until cells were completely lysed. Lysates were centrifuged at 12,000×g for 30 min at 4 °C. Finally, the fusion protein was purified using His-Bind columns (Qiagen, Venlo, The Netherlands) and analyzed by SDS-PAGE using gels containing 12% polyacrylamide. After electrophoresis, the gel was transferred to a nitrocellulose membrane (Amersham, Bucks, UK), and the nitrocellulose membranes were blocked for 4 h with TBST solution containing 5% skim milk powder. After being washed with TBS five times, the blots were incubated with an anti-His6-tag monoclonal antibody (Novagen, Darmstadt, Germany) at 37 °C for 2 h. After the reaction, the membranes were washed for three times with phosphate buffer saline (PBS) solution (pH=7.4) for 10 min each time. The membranes were incubated for 2 h at 37 °C with horseradish peroxidase (HRP)-conjugated goat anti-mouse IgG in TBST buffer (Tris 10 mM, NaCl 150 mM, Tween-20 0.05%(V/V), HCl to PH 7.5) containing 5% skim milk. At the end of hybridization, the color was developed with TMB (promega). When the band appeared, the film was washed with water immediately to stop the reaction.

### Determination of nitrite reductase activity

*E. coli* BL21 containing the recombinant plasmid *nirS*-pET28a or pET28a-*nirS*-EGFP was inoculated into 50 ml of LB medium and cultured at 37 °C and 200 rpm for 8 h until the OD value of the culture was about 0.5. The BL21 strain was induced at a concentration of 0.2 mM IPTG at 16 °C and 110rpm for 16h. The induced bacterial liquid was broken by sonication and centrifuged at 8000×g. The obtained supernatant was a crude enzyme sample of nitrite reductase. The crude enzyme (100 μl) was added into the reaction system containing Phosphate buffer (pH 7.1) 10 mM, sodium nitrite NaNO_2_ 1 mM, methyl amethyst 1 mM, sodium thiosulfate 5 mM, and ddH_2_O up to 2 ml in a water bath at 30 °C for 30 min before the reaction was terminated by violent shaking. Griess color developer was then added and reacted at room temperature for 20 min; the absorbance was measured at 420 nm with an ultraviolet spectrophotometer (Lambda 25, PerkinElmer Company, USA).

### Isolation of OMVs

OMVs were isolated from late log-phase culture of *E. coli*-*nirS* followed by J. Habier et al. (45). In brief, 500 ml of LB medium was inoculated with 5 ml of culture grown to OD_600_ equal to 1.0 and was incubated at 37 °C on a shaker at 150 rpm in 50mg l^-1^ kanamycin. To isolate OMVs from cultures grown under heavy metal stress, liquid cultures were grown on a shaker with sub-inhibitory concentrations of copper ion (25 mg l^-1^ or 50 mg l^-1^) or chromium ion (5 mg l^-1^ or 25 mg l^-1^). For temperature stress experiments, the temperature was raised to 37 °C. After cells were centrifuged using a 50 ml tube at 8000×g for 25 min, the supernatant was filtered through a 0.45 mm membrane (Millipore Corporation, Bedford, MA, USA) to remove cells. The filtrate was concentrated 50-fold using 100 kDa ultrafiltration tube (Millipore Corporation, Bedford, MA, USA) and then filtered through a 0.22 μm vacuum membrane to remove any remaining cells and cell debris. The filtrate was put into an ultra-high-speed centrifuge (Optima MAX-TL, Beckman, USA) at 160,000×g and 4°C for 3 h. The pellet of the extracted OMVs suspended in PBS was ultracentrifuged at 160,000×g and 4 °C for 1 h for 2 times. The pellet of the extracted OMVs was finally re-suspended in 500 μl of PBS.

### Purification of OMVs

Crude OMVs were purified by density gradient as described by H. Chutkan et al. (46). Briefly, 600 μl of crude OMVs was added to the bottom of a 5 ml centrifuge tube, and 400 μl of 60%, 1 ml of 40%, 1 ml of 35%, 1 ml of 30% and 1 ml of 25% OptiPrep/PBS gradient centrifuge solutions were added from bottom to top. The gradients were centrifuged (160,000×g) for 6 h at 4 °C. Five fractions of equal volume were collected from the bottom. Each fraction was washed at the same speed by adding PBS buffer 10 times and subjected to 10% SDS-PAGE. Protein content of purified OMVs was estimated using the Bradford assay (47), which can also be used for quantitative calculation of vesicles.

### Determination of aerobic denitrification gene (nirS) and total DNA in OMVs

The total DNA in OMVs was quantified by an ultramicro spectrophotometer (NanoDrop 2000C, Thermo Fisher, USA). Briefly, vesicular proteins of 50 μg were treated with 2 μl proteinase K (20 mg ml^-1^) at 37 °C for 30 min to hydrolyze the bacteriophage. Proteinase K was inactivated at 80 °C for 5 min. The surface-associated DNA of OMVs was degraded by adding 2 μl of DNase I (5 U μl^-1^) at 37 °C for 1 h. DNase I was permanently inactivated at 80 °C for 5 min by adding 0.5 M EDTA (Sagon Biotech, Shanghai, China). Subsequently, treated OMVs were lysed with 0.25% Triton X-100 solution (Sagon Biotech, Shanghai, China) for 30 min at 37 °C. DNA released by OMVs was extracted by a SanPrep column PCR product purification kit (Sagon Biotech, Shanghai, China) and quantified by an ultramicro spectrophotometer. To determine the *nirS* gene existed in OMVs, extracted DNA was re-suspended in 15 μl of TE buffer DNA. DNA of OMVs from *E. coli*-*nirS* was amplified by PCR using primers (Table S1). PCR products were analyzed by gel electrophoresis on 1% agarose.

### Transmission electron microscopy (TEM)

Vesicle suspension from *E. coli*-*nirS* was negatively stained with freshly prepared 2% aqueous uranyl acetate on carbon-coated nickel grids (300 mesh) (Sigma). The excess stain was blotted, and the grid was washed once with distilled water and dried. Micrographs were obtained by TEM (JEM1400, Japan). To observe the cellular structure and OMVs release from *E. coli*, the bacterial culture was pipetted into a 1.5 ml centrifuge tube and centrifuged at 2000 rpm for 10 min. Then, the supernatant was removed, and cells were washed with PBS solution (pH=7.4) for 3 times. Afterward, the collected bacterial samples were fixed with 2.5% cold glutaraldehyde for overnight at 4 °C. After being washed, the bacterial sample was dehydrated by ethanol and acetone, embedded and stained with Osmium tetroxide (OsO4). The treated sample was heated at 70 °C overnight using a heating polymerization apparatus. Ultra-thin slices (50–70 nm) of the sample were made by an ultra-thin slicing machine. Finally, the section was placed on carbon-coated nickel grids and stained with freshly prepared 2% aqueous uranyl acetate for 3 min, after which *E. coli* was observed under TEM.

### Transformation of aerobic denitrification gene (nirS) via OMVs in liquid cultures

*E. coli* BL21 cells were collected (2000×g) at 4 °C for 10 min in a logarithmic growth period. After removing the supernatant, cells were re-suspended by 100 μl of fresh SOC medium (59% Tryptone; 15% Yeast Extract; 1.4% NaCl; 15% MgSO_4_·7H_2_O; 11% D-Glucose) and then mixed with different amounts of purified OMVs (containing 50, 100 and 200 μg of protein) released from *E. coli*-*nirS*. The suspensions were incubated statically at 37 °C for 4 h and then for another 4 h on a shaker at 150 rpm. LB broth (10 ml) was then added to the suspension, and the culture continued at 37 °C overnight at 150 rpm. Finally, the overnight culture was centrifuged at 8000×g and then suspended in 500 μl LB medium. The 250 μl resuspension was placed on a plate with LA medium and 50 mg l^-1^ kanamycin. Successful transformation in *E. coli* BL21 was confirmed by observing positive colony numbers and the *nirS* gene by PCR. To ensure that the transformation experiment was mediated by DNA in the OMVs, the OMVs in the experiment have been treated with DNase I and proteinase K. Gene transfer frequencies were calculated from three independent experiments as the number of gene transfer events over the number of recipient cells.

### Laser confocal scanning microscope

The purified OMVs were stained with CellTracker™ CM-DiI Dye (Thermo Fisher), which is non-fluorescent in aqueous solution, and were labeled by binding OMVs to lipid molecules of the membrane structure. OMVs were incubated with 0.5 μM CM-DiI dye at 37 °C for 30 min and then centrifuged at 20,000 × g and 4°C for 2 h. The stained OMVs were washed twice with PBS buffer (pH 7.4) and then re-suspended in 100 μl PBS. *E. coli* BL21 cells were stained with VybrantTM DiO live cell tracer (ThermoFisher). Cell-labeling solution (5 μl) was put into 1 ml of *E. coli* BL21 cell suspension (1×10^6^ ml^-1^) and incubated at 37 °C for 20 min. Finally, the cell suspension was centrifuged at 2000 × g for 10 min. After the supernatant was removed, cells were washed twice with PBS buffer (pH 7.4) and then re-suspended in 500 μl PBS before stained OMVs with CM-DiI fluorescence were added. Then, the mixed liquid was incubated statically for 45 min at 37 °C. OMVs and *E. coli* BL21 were separately analyzed using an excitation wavelength of 553 nm and 484 nm under a laser confocal scanning microscope (LSM710, Zeiss, Germany) within 1 h. *E. coli* BL21 successfully transformed with pET28a-*nirS*-EGFP recombinant plasmid via OMVs was induced with 0.2 mM IPTG for 12 h at 16 °C and observed at 488 nm excitation wavelength by LCSM.

## Acknowledgments

This study was supported by a project funded by the Priority Academic Program Development of Jiangsu Higher Education Institutions (PAPD).

## Author contributions

The conception or design of the study, Y. Luo, J. Miao, W. Qiao; The acquisition, Y. Luo; Analysis, Y. Luo, J. Miao; Interpretation of the data, Y. Luo, J. Miao; Writing – Original Draft, Y. Luo; Writing – Review and Editing, W. Qiao; Funding Acquisition, W. Qiao

